# Choosing representative proteins based on splicing structure similarity improves the accuracy of gene tree reconstruction

**DOI:** 10.1101/2020.04.09.034785

**Authors:** Esaie Kuitche Kamela, Marie Degen, Shengrui Wang, Aïda Ouangraoua

**Affiliations:** Department of Computer Sicence, University of Sherbrooke, Quebec, J1K 2R1, Canada

**Keywords:** Gene tree, Phylogenetics, Algorithms, Alternative splicing, Splicing homology, Fuzzy C-means

## Abstract

Constructing accurate gene trees is important, as gene trees play a key role in several biological studies, such as species tree reconstruction, gene functional analysis and gene family evolution studies. The accuracy of these studies is dependent on the accuracy of the input gene trees. Although several methods have been developed for improving the construction and the correction of gene trees by making use of the relationship with a species tree in addition to multiple sequence alignment, there is still a large room for improvement on the accuracy of gene trees and the computing time. In particular, accounting for alternative splicing that allows eukaryote genes to produce multiple transcripts/proteins per gene is a way to improve the quality of multiple sequence alignments used by gene tree reconstruction methods. Current methods for gene tree reconstruction usually make use of a set of transcripts composed of one representative transcript per gene, to generate multiple sequence alignments which are then used to estimate gene trees. Thus, the accuracy of the estimated gene tree depends on the choice of the representative transcripts. In this work, we present an alternative-splicing-aware method called Splicing Homology Transcript (SHT) method to estimate gene trees based on wisely selecting an accurate set of homologous transcripts to represent the genes of a gene family. We introduce a new similarity measure between transcripts for quantifying the level of homology between transcripts by combining a splicing structure-based similarity score with a sequence-based similarity score. We present a new method to cluster transcripts into a set of splicing homology groups based on the new similarity measure. The method is applied to reconstruct gene trees of the Ensembl database gene families, and a comparison with current EnsemblCompara gene trees is performed. The results show that the new approach improves gene tree accuracy thanks to the use of the new similarity measure between transcripts. An implementation of the method as well as the data used and generated in this work are available at https://github.com/UdeS-CoBIUS/SplicingHomologGeneTree/.

## 1 Introduction

In the last two decades, several phylogeny methods based on multiple alignment of homologous sequences have been developed and used for gene tree reconstruction (eg. PhyML (Guindon and Gascuel 2003), RAxML (Stamatakis 2006), PhyloBayes (Lartillot and Philippe 2004)). It has been shown that the gene trees reconstructed solely based on multiple sequence alignments often contain many errors and uncertainties, due to numerous reasons which include the quality of gene annotations, the quality of multiple sequence alignments, the quantity of substitutions in the alignments, and the accuracy of the phylogeny reconstruction methods (Hahn 2007; Rasmussen and Kellis 2007). To overcome these limitations, the development of phylogenomics methods which make use a known species tree in addition to multiple sequence alignments has been fruitful to improve the accuracy of reconstructed gene trees (eg. TreeBeST (Schreiber et al. 2014), NOTUNG (Chen, Durand, and Farach-Colton 2000), ALE (Szöllősi et al. 2013), GSR (Åkerborg et al. 2009), TERA (Scornavacca, Jacox, and Szöllősi 2015), SPIMAP (Rasmussen and Kellis 2011), GIGA (Thomas 2010)). Several methods for gene tree correction guided by species tree have also been developed (eg. ProfileNJ (Noutahi et al. 2016), TreeFix (Wu et al. 2013), TreeFix-DTL (Bansal et al. 2015), LabelGTC (El-Mabrouk and Ouangraoua 2017), Refine-Tree(Lafond et al. 2013), and (Górecki and Eulenstein 2012)). Another category of methods aims at jointly inferring gene trees and species trees (eg. PHYLDOG (Boussau et al. 2013), DLRS (Sjöstrand et al. 2012), PhyloNet (Y. Wang and Nakhleh 2018), BP&P (Rannala and Yang 2017)). The later provide accurate reconstructions by modeling and resolving the discordance between gene trees and species trees (Maddison 1997), but they are also limited by a scalability problem for larger data sets. Despite the improvement in the accuracy and resolution of gene trees reconstructed by all these approaches, there is still a large room for improvement on the accuracy of gene trees and the computing time.

In this work, we approach the question of improving the quality of multiple sequence alignments used in phylogenomics methods for gene tree reconstruction, by choosing wisely the set of proteins representing genes of a gene family. A key factor for the accuracy of multiple sequence alignment is that aligned sequences should be homologs. It is well known that alternative splicing (AS) is a ubiquitous process in eukaryote organisms by which multiple transcripts and proteins are produced from a single gene (Keren, Lev-Maor, and Ast 2010; Nilsen and Graveley 2010; Blencowe 2006; Blencowe 2017; Modrek and Lee 2002; Kuitche, Lafond, and Ouangraoua 2017; Iñiguez and Hernández 2017). Gene tree reconstruction methods account for this information, by selecting and making use of one reference transcript/protein sequence per gene to reconstruct the gene trees that populate gene tree databases (eg. EnsemblCompra (Vilella et al. 2009), PhylomeDB (Huerta-Cepas et al. 2014), Panther (Mi, Muruganujan, and Thomas 2012), TreeFam (Schreiber et al. 2014)). Among existing gene tree databases, the EnsemblCompara method is the most popular one (Vilella et al. 2009). Like most of the current gene tree reconstruction methods, the EnsemblCompara method chooses the longest transcript for each gene as its representative transcript to generate a multiple sequence alignment. This choice ignores the transcripts of all the other genes of the gene family. The rationale of this choice is that the longest transcript is the more representative because its coverage of the gene is the largest (Kasukawa et al. 2004). However, there is no guarantee that the set of representative transcripts composed of the longest transcript of each gene constitutes the set of the most homologous transcripts between genes. The homology level of two transcripts from two homologous genes can be measured by the quantity of homologous nucleotides and exons shared by the transcripts. The longest transcripts of two homologous genes may be composed of completely different exons. Thus, choosing the longest transcript for each gene, independently of other genes, may lead to poorly homologous transcripts, and then erroneous sequence alignments and gene trees. One of the main consequences of choosing a set of non homologous transcripts is that the alignment of those sequences will forcibly align some of those sequences together, which will muddle the phylogenetic signal. Ideally, the set of transcripts chosen to estimate a gene tree must be equivalogs (Haft et al. 2001). Equivalogs are homologous transcripts that have conserved their functions since their last common ancestor. Equivalogs share the same functions, as well as conserved sequences and homologous exons, which results in similar splicing structures.

Classical methods for clustering homologous transcripts into equivalog families and automated functional identification of proteins rely on sequence and folding structure similarity. This approach ignores the splicing structure similarity between transcripts (Haft et al. 2001; Pandit et al. 2002; Zhang and Skolnick 2005; Balamurugan, Dekker, and Waldmann 2005; Balaji and Srinivasan 2007; Xu and Zhang 2010). However, it has been shown that the splicing structure of homologous sequences is often more conserved than the nucleotide sequences (Betts et al. 2001; Abril, Castelo, and Guigó 2005; Poulos et al. 2011). It has also been shown that the splicing structure contains information that can improve protein and transcript comparison (Abascal, Tress, and Valencia 2015; Abascal, Ezkurdia, et al. 2015). IsoSel (Philippon et al. 2017) is a tool devoted to the selection of equivalogs in the context of phylogenetic reconstruction. It provides a better alternative of choosing the longest isoforms per gene. It uses multiple sequence alignment and bootstrapping to identify equivalogs.

In this paper, we introduce a new framework for the reconstruction of eukaryote gene trees. We consider all transcripts of all genes of a gene family, and we define a new similarity measure by combining splicing structure-based similarity and sequence-based similarity. Based on the all-against-all similarity matrix between transcripts, we propose a clustering method, based on an improved version of the fuzzy c-means (FCM) algorithm (Bezdek, Ehrlich, and Full 1984) and a latent representation of transcripts. This method allows to obtain splicing homology groups, that represent estimated equivalog families. Finally, we use a multiple sequence alignment of the selected representative transcripts to build the gene tree.

The paper is organized as follows: the next section describes the method devised to estimate gene trees by taking into account the splicing structure of transcripts. The Results and Discussion section presents the results of a comparative analysis between the gene trees estimated using the new method, IsoSel, and, the EnsemblCompara gene trees which are currently the most largely used. The trees reconstructed with our method compare favorably with the trees from the EnsemblCompara database, in terms of the significance of a likelihood difference computed using an Approximately unbiased test (Shimodaira 2002), and in terms of the reconciliation with a species tree.

## 2 Materials and Methods

We first introduce some formal definitions that will be useful for the description of the method. Next, we present the definition of the new measure of similarity between transcripts, and then the new algorithm for clustering transcripts from a similarity matrix. In the last part of this section, we present how the gene trees are estimated from a set of representative transcripts. An overview of the method is depicted in Figure 1.

**Fig. 1:**
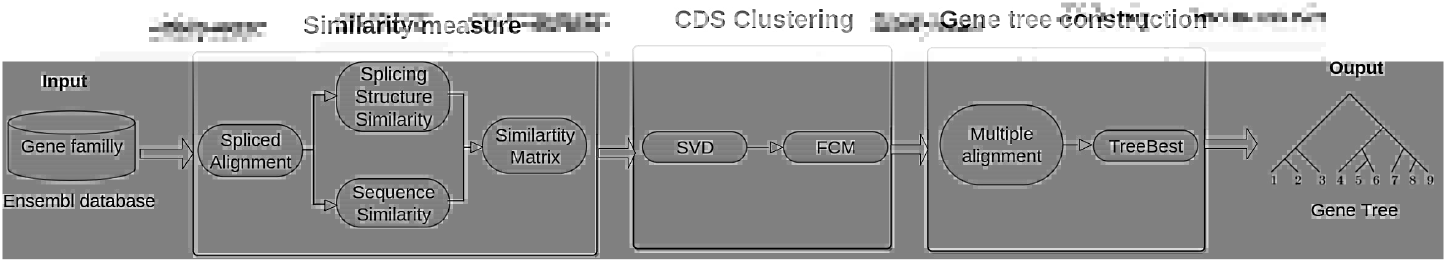
Overall view of the method: The first step consists in the computation of the similarity matrix between Coding DNA Sequence (CDS) of transcripts. The second step consists in grouping CDS into splicing homology groups and, finally, the last step consists in aligning the representative CDS and building the gene tree.

### 2.1 Preliminary definitions

#### Gene and CDS

A gene sequence is a DNA sequence on the alphabet of nucleotides *σ* = {*A, C, G, T*}. Given a gene sequence *g*, an exon of *g* is a transcribed segment of *g*. A coding DNA sequence (CDS) of *g* is a concatenation of translated exons of *g*. The following is an example of toy gene *g* with two alternative CDS *c*_1_ and *c*_2_. The alignment of *g*, *c*_1_ and *c*_2_ shows that *c*_1_ is composed of 3 exons, and *c*_2_ of 4 exons. Here, CDS refers to transcript or isoform.

~~~
>g
CGTATGGAATGCGTAAGAAGCAGGTCGTAAGCATACGTGGGTAAGGGGAATGATTGAAAG
>c1
ATGCAAGCAGGTCGGGAATGA
>c2
ATGGAATGCAAGCAGCATACGTGGGGGAATGATTGA
~~~

~~~
g : CGTATGGAATGCGTAAGAAGCAGGTCGTAAGCATACGTGGGTAAGGGGAATGATTGAAAG
c1: --------ATGC-----AAGCAGGTC-------------------GGGAATGA
c2: ---ATGGAATGC-----AAGCAG--------CATACGTGG-----GGGAATGATTGA
~~~

#### Multiple splicing structure

Given a set of CDS 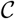 from a set of homologous genes, a segment class of 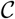 is a set composed of exactly one, possibly empty, segment from each element of 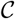, such that these segments are homologous, i.e descending from the same ancestral segment. A segment class must contain at least one non-empty segment. A multiple splicing structure of a set of CDS 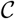 is a chain *E* = {*E*[1]*, …, E*[*n*]} of segment classes of 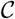 in which no pair of classes *E*[*i*] and *E*[*j*] contains an overlapping segment, and each CDS *c* in 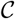 is the concatenation of its segments belonging to classes of *E*. Figure 2 presents an example of multiple spliced structure of three CDS decomposed into three segment classes. In practice, a set of homologous CDS is decomposed into a chain of segment classes corresponding to homologous exon classes.

**Fig. 2:**
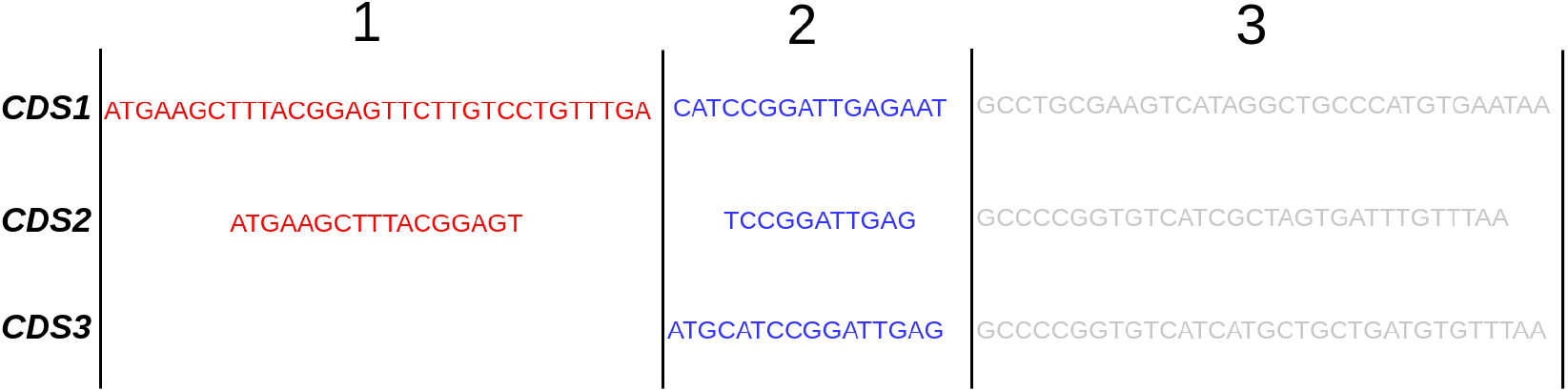
Multiple splicing structure: Multiple splicing structure of three homologous CDS decomposed into three segment classes represented by different colors: red color for the first class, blue for the second class, and grey for the third class. Each CDS has a non-empty segment in all segment classes, except CDS3 which has an empty segment in the first class.

#### A multiple spliced alignment

Given a set of homologous CDS 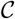 decomposed into a multiple splicing structure *E* = {*E*[1]*, …, E*[*n*]}, the multiple spliced alignment of 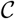 corresponding to *E* is a chain *A* = {*A*[1], …, *A*[*n*]} such that each *A*[*i*] is an alignment of the segment class *E*[*i*]. For any CDS *c* in 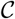, *A*_*c*_ = {*A*[*i*]_|*c*_|1 ≤ *i* ≤ *n*} is the chain of *n* gapped segments of *c* induced by the multiple spliced alignment *A*. Given a multiple spliced alignment *A*, |*A*| denotes the number of sub-alignments composing *A*, i.e. the number of segment classes in the corresponding multiple spliced alignment. Given a gapped nucleotide segment *x*, *len*(*x*) denotes the number of nucleotides in *x*. Figure 3 gives an example of multiple spliced alignment corresponding to the multiple splicing structure of three CDS from Figure 2.

**Fig. 3:**
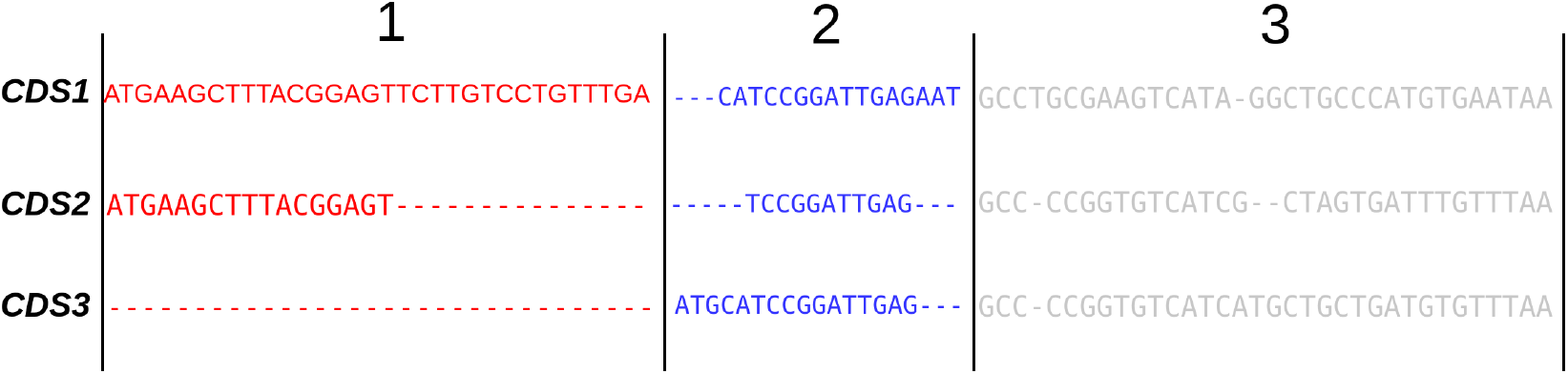
Multiple spliced alignment: a multiple spliced alignment *A* corresponding to the multiple splicing structure of three CDS from Figure 2. For instance, |*A*| = 3, *len*(*A*[1]_|*CDS*3_) = 0, and *len*(*A*[2]_|*CDS*3_) = 11.

### 2.2 Similarity measure

The similarity measure between two CDS, *x* and *y*, of a set of homologous CDS is defined as follows:

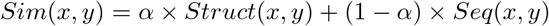

*Struct*(*x, y*) is a splicing structure similarity between *x* and *y*, *Seq*(*x, y*) is a sequence similarity between *x* and *y*, and *α* is such that 0 ≤ *α* ≤ 1 and defines the relative weights of *Struct*(*x, y*) and *Seq*(*x, y*) in the similarity score.

The value of *Struct*(*x, y*) is defined by combining three similarity scores:

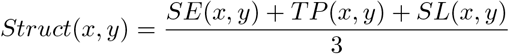

*SE*(*x, y*), *TP* (*x, y*) and *SL*(*x, y*) are respectively a splicing event (SE) similarity score, a translation phase (TP) similarity score, and a segment length (SL) similarity score between *x* and *y*, defined thereafter. Here, translation phase refers to the reading frame. It may take the values 0, 1 or 2.

#### Splicing event similarity

Given a multiple spliced alignment *A* of a set of CDS 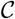, we consider four possible types of alternative splicing events for each gapped segment *A*[*i*]_|*x*_ such that 1 ≤ *i* ≤ |*A*| and 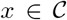. Figure 4 illustrates the different types of alternative splicing events. An exon skipping (ES) arises when the segment contains only gaps. For instance, the first segment of CDS3 in Figure 4 has an ES event. A 3’ alternative splice site (3P) event arises when the segment starts with gaps. For instance, the second segments of CDS2 and CDS3 have a 3P event. A 5’ alternative splice site (5P) event arises when the segment ends with gaps. For instance, the first segment of CDS2 as well as the second segments of CDS1 and CDS3 have a 5P event. An internal gap (IG) event arises when there is a set of consecutive gaps within the segment. For instance, the third segment of CDS1, CDS2 and CDS3 have at least one IG event.

**Fig. 4:**
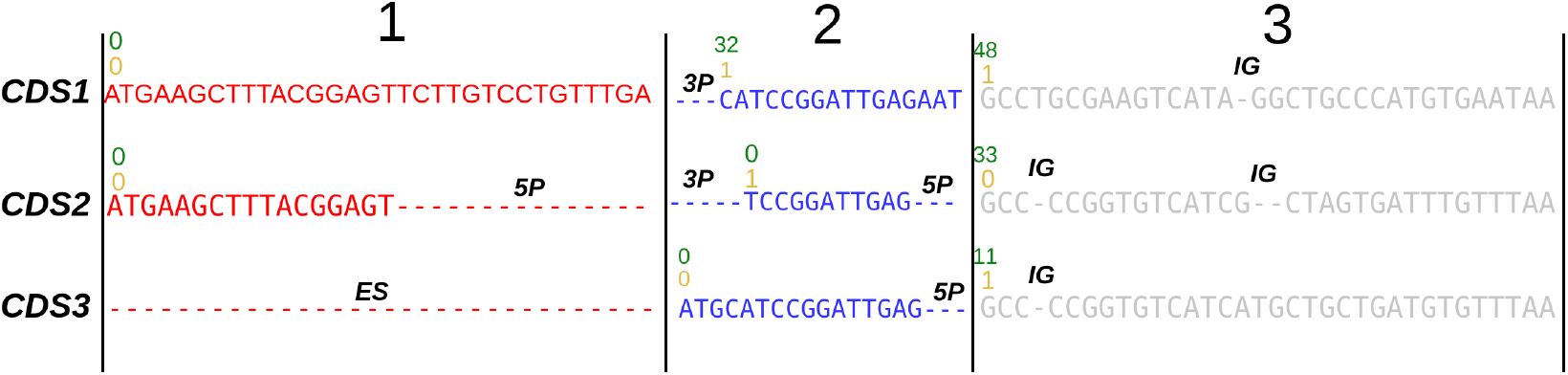
Alternative splicing events: The multiple spliced alignment of Figure 3 with gapped segments annotated with their alternative splicing events, translation phases, and segment lengths. Above the first nucleotide of each segment, the number in green represents the position of the first nucleotide in the CDS and the number in orange represents the translation phase of the exon. The alternative splicing events of each segment are indicated (Exon Skipping(ES), 3 Prime Extension(3P), 5 Prime Extension(5P), or Internal Gap(IG)).

For a CDS *x*, we denote by *e*[*i*]_|*x*_ the set of alternative splicing events displayed by the gapped segment *A*[*i*]_|*x*_. Given two CDS *x* and *y* from 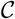, the splicing event similarity between the two gapped segments *A*[*i*]_|*x*_ and *A*[*i*]_|*y*_ is the ratio of the number of common events in the two sets *e*[*i*]_|*x*_ and *e*[*i*]_|*y*_:

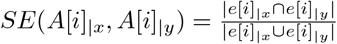

The splicing event similarity between *x* and *y* is then given by:

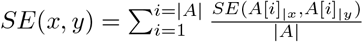

#### Translation phase similarity

Given a multiple spliced alignment *A* of a set of CDS 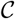, for each gapped segment *A*[*i*]_|*x*_ such that 1 ≤ *i* ≤ |*A*| and 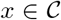, we denote by *phase*[*i*]_|*x*_ the translation phase of the segment *A*[*i*]_|*x*_. The value of *phase*[*i*]_|*x*_ equals the position of the first nucleotide of *A*[*i*]_|*x*_ in the sequence *x* modulo 3. Given two CDS *x* and *y* from 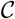, the translation phase similarity between the two gapped segments *A*[*i*]_|*x*_ and *A*[*i*]_|*y*_ is 1 if the two segments have the same translation phase, and 0 otherwise:

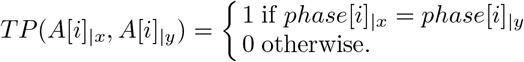

The translation phase similarity between *x* and *y* is then given by:

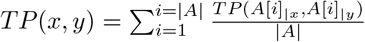

#### Segment length similarity

Given two CDS *x* and *y* from 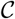, the segment length similarity between the two gapped segments *A*[*i*]_|*x*_ and *A*[*i*]_|*y*_ is given by:

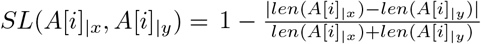

The segment length similarity between *x* and *y* is then given by:

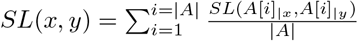

#### Sequence similarity

Giving a multiple spliced alignment *A* of a set of CDS 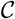, we use the substitution matrix *BLOSUM100* as substitution model to compute the sequence similarity score between each pair of CDS in 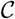.

Based on the definition of the similarity score *Sim*(*x, y*) between two CDS *x* and *y*, an all-against-all similarity matrix is computed for the set of homologous CDS 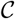 of any gene family.

### 2.3 CDS Clustering

In order to determine the representative CDS of a set of homologous genes, CDS are first clustered into splicing homology groups. It has been demonstrated that, for clustering problems where clusters may overlap each other, fuzzy clustering methods, which can assign a data to more than one cluster, achieve higher performance than hard clustering methods, which assign each data to a single cluster (Sun, S. Wang, and Jiang 2004). The CDS clustering problem is best approach as a fuzzy clustering problem because a CDS in one gene may be similar in sequence and splicing structure to several alternative CDS of another gene, and splicing homology groups could overlap.

The Fuzzy C-means (FCM) algorithm and its derivatives are the most widely used fuzzy clustering algorithm (Bezdek, Ehrlich, and Full 1984). As input, they require a representation of the data, generally a vectorial representation, which facilitates computing of the centroid of a cluster of data. They also require as input the number of clusters in the given data set. The clustering method starts with a transformation of the CDS representation into a vectorial representation, and the application of the FCM-Based Splitting Algorithm (FBSA) (Sun, S. Wang, and Jiang 2004) to cluster CDS.

#### Vectorial representation of CDS

In this section, we propose to compute a latent vectorial representation of CDS from a *N × N* similarity matrix *S*, containing the pairwise similarity scores between the *N* CDS of a set of homologous genes. Such a representation allows CDS to be represented in a new geometric space while preserving their mutual similarity and facilitating computing of centroids in clustering algorithms. Each CDS will be represented as an N-dimensional vector by applying a spectral decomposition on the similarity matrix *S*. The spectral decomposition of a symmetric *N* × *N* matrix *S* is the decomposition of *S* into *S* = *UDU*^*T*^, such that *U* is an orthogonal *N* × *N* matrix and *D* is a *N* × *N* diagonal matrix. The columns of *U* correspond to the eigen vectors of *S*, and the diagonal entries of *D* correspond to the eigen values.

We define an *N* × *N* matrix 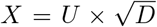, such that each row *x*_*i*_ of *X* is the new vectorial representation of the i^*th*^ CDS from the similarity matrix *S*. This transformation is valid because the similarity score *S*_*ij*_ between the i^*th*^ and j^*th*^ CDS is now approximately equal to the inner vector product *< x*_*i*_, *x*_*j*_ *>* of the vectors representing the CDS. Thus, the relative similarity scores between CDS are preserved by the transformation, and the vectors {*x*_1_, …, *x*_*N*_} can be clustered using a fuzzy clustering algorithm.

#### Application of the FBSA for fuzzy clustering

The FCM algorithm requires, as input parameter, the number of clusters in the data-set in order to be clustered (Bezdek, Ehrlich, and Full 1984). Given *N* data vectors *X* = {*x*_1_, *x*_2_, …, *x*_*N*_} to cluster, and *c* the expected number of clusters, the FCM algorithm searches for a *N* × *c* non-negative matrix *M* called the fuzzy partition matrix. Each value *m*_*ki*_ such that 1 ≤ *k* ≤ *N* and 1 ≤ *i* ≤ *c* in the matrix *M* is the membership value of vector *x*_*k*_ to the i^*th*^ cluster, and for any vector 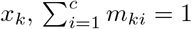. Given a *N* × *c* membership matrix *M*, the centre of each cluster *i*, 1 ≤ *i* ≤ *c*, is computed as the vector:

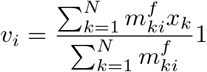

The exponent *f >* 1 in the cluster centre formula is a parameter called a fuzzifier. Given a set of cluster centres *V* = {*v*_1_, …, *v*_*c*_}, the values of the *N× c* membership matrix *M* are computed as follows:

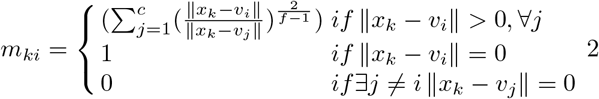

The formula (1) and (2) are derived from minimization of the following objective function *F*_*f*_.

Given *N* data vectors *X* = {*x*_1_, *x*_2_, …, *x*_*N*_} and an expected number of clusters *c*, the FCM algorithm computes a membership matrix *M* and a set of cluster centres *V* = {*v*_1_, …, *v*_*c*_} that minimizes the following function

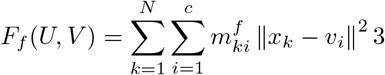

The FCM algorithm starts by randomly initializing the cluster centres *V* = {*v*_1_, …, *v*_*c*_}. Then it iteratively recalculates the membership matrix *M* using equation (2) and a new set of cluster centres using equation (1) until a set of stable cluster centres is reached.

The main limitation of the FCM algorithm is that it requires the number of clusters as input. Several derivatives of the FCM algorithm have been developed for automatically determining the number of clusters. As input parameters, they take a lower bound *c*_*min*_ and an upper bound *c*_*max*_ for the number of clusters, compute an optimal clustering using FCM for each number of cluster *c* such that *c*_*min*_ ≤ *c* ≤ *c*_*max*_, and choose the best value of *c* based on a cluster validity criterion.

The FBSA algorithm is a derivative of the FCM algorithm that allows to cluster a data set while automatically determining the number of clusters (Sun, S. Wang, and Jiang 2004). FBSA improves on other derivatives of the FCM algorithm to allows automated selections of the number of clusters. It reduces the randomness in the initialization of cluster centres at the beginning of each clustering phase. For each clustering phase with a value of *c* such that *c*_*min*_ < *c* ≤ *c*_*max*_, the clustering is carried out starting with the previously obtained *c* – 1 clusters and splitting the worst cluster in two clusters to obtain *c* clusters. The worst cluster is the one corresponding to the minimum value of a score function *S*(*i*) associated with each cluster *i* as follows:

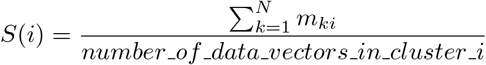

A second improvement of FBSA in the selection of the best number of clusters is the use of a new validity index *V*_*d*_(*c*) based on a linear combination of cluster compactness and separation that is efficient even when clusters overlap each other. The validity index *V*_*d*_(*c*) is computed for each cluster number *c* to choose the number of clusters that maximizes *V*_*d*_(*c*). The detailed description of FBSA and the definition of the validity index are provided in (Sun, S. Wang, and Jiang 2004).

Given a *N × N* similarity matrix *S* on a set of *N* homologous CDS 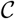 of a gene family, we extend the FBSA method to cluster 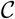 into subsets of splicing homology groups, and select a set of representative CDS as follows:

1. Apply a spectral decomposition on the matrix *S* to obtain a vectorial representation *X* = {*x*_1_, …, *x*_*N*_} of the *N* CDS in 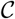;
2. Apply FBSA on *X* = {*x*_1_, …, *x*_*N*_} and choose the best clustering corresponding to a number *c*_*opt*_ of clusters maximizing *V*_*d*_(*c*_*opt*_);
3. Select the cluster *i* in the *c*_*opt*_-clustering such that *S*(*i*) is maximal.
4. For each gene *g*, select as representative CDS, the CDS *k* of *g* such that the membership *m*_*ki*_ of *x*_*k*_ to the i^*th*^ cluster is maximum.

### 2.4 Gene tree construction

In order to carry out a comparative study on the effect of the choice of representative CDS for gene tree construction, the new method and IsoSel method use the same steps as the Ensembl method, except for the representative CDS selection step. After selecting the representative CDS, the third methods compute a multiple sequence alignment of the protein sequences corresponding to the representative CDS using Mafft (Katoh and Standley 2013). Next, the CDS back-translated protein alignment and the species tree are used to estimate the gene trees with the TreeBest phylogenetic reconstruction method. TreeBest is a method for building gene trees using a known species tree (Schreiber et al. 2014).

## 3 Results and Discussion

In order to evaluate the new gene tree reconstruction method, we compare it with the method from EnsemblCompara and the method from IsoSel, based on a dataset of gene families from the EnsemblCompara database (Vilella et al. 2009). In the remaining section, since the differences between the three compared methods lies within the approach used to select a set of representative transcripts for genes, the methods are named according to the latter. The EnsemblCompara method that selects the longest transcript of each gene is named the Longest Transcript (`LT`) method, The IsoSel method that select a set of isoform transcript is name Isosel, and the new method that computes and uses a splicing homology group is called the Splicing Homology Transcript (`SHT`) method.

### 3.1 Dataset and compared methods

The initial dataset contains 2036 gene families of 173 species selected randomly from the EnsemblCompara database (release 98) (ibid.). The data for each gene family is composed of a set of homologous genes, their Coding Dna Sequences (CDS) and the gene tree from the EnsemblCompara database called the LT_db gene tree. For each gene, only CDS that start with a start codon and whose lengths are multiple of 3 are considered, because the information on the coding phase is required for computing the similarity scores between CDS in the first step of the method, and the translations into protein sequences are needed for building the gene tree using TreeBest (Schreiber et al. 2014) in the last step of the method. Thus, the LT_db gene tree contains only genes for which at least one CDS is considered. The LT_db gene tree is induced from the EnsemblCompara database gene tree that is often on a larger set of genes. IsoSel gene tree is the tree obtained by applying TreeBest on the transcript isoforms return by IsoSel.

Figure 5 shows some statistics on the number of genes per family and the number of CDS per gene in the dataset. Figure 5 (a) shows the percentage of gene families per ratio of the number of genes retained in this study over the number of genes in the EnsemblCompara database. For instance, we observe that in more than 45% of gene families, the number of genes used in this study is less than 10% of the number of genes in the EnsemblCompara gene family. Figure 5 (b) shows the percentage of genes per number of CDS. We observe that almost 80% of genes have only one CDS, and 15% of genes have two CDS. Figure 5 (c) shows the percentage of gene families per percentage of genes familly having a single CDS. We note that more than 60% of gene families have more than 90% of genes having a single CDS. Out of 2036 gene families, 1313 families have a single CDS per gene, which induces a single possible set of representative transcripts, thus the same tree is reconstructed using the two methods. These families were discarded, leaving a dataset with 723 gene families for the comparison. We call this dataset the 723-dataset, and the initial dataset is called the 2036-dataset. Figure 5 (d) and (e) show the statistics on the number of CDS per gene in the 723-dataset. For instance, we observe that the number of single-CDS genes in the dataset is still high (more than 55 %), but the percentage of gene families that have more than 90% single-CDS genes is lower (less than 5% of gene families).

**Fig. 5:**
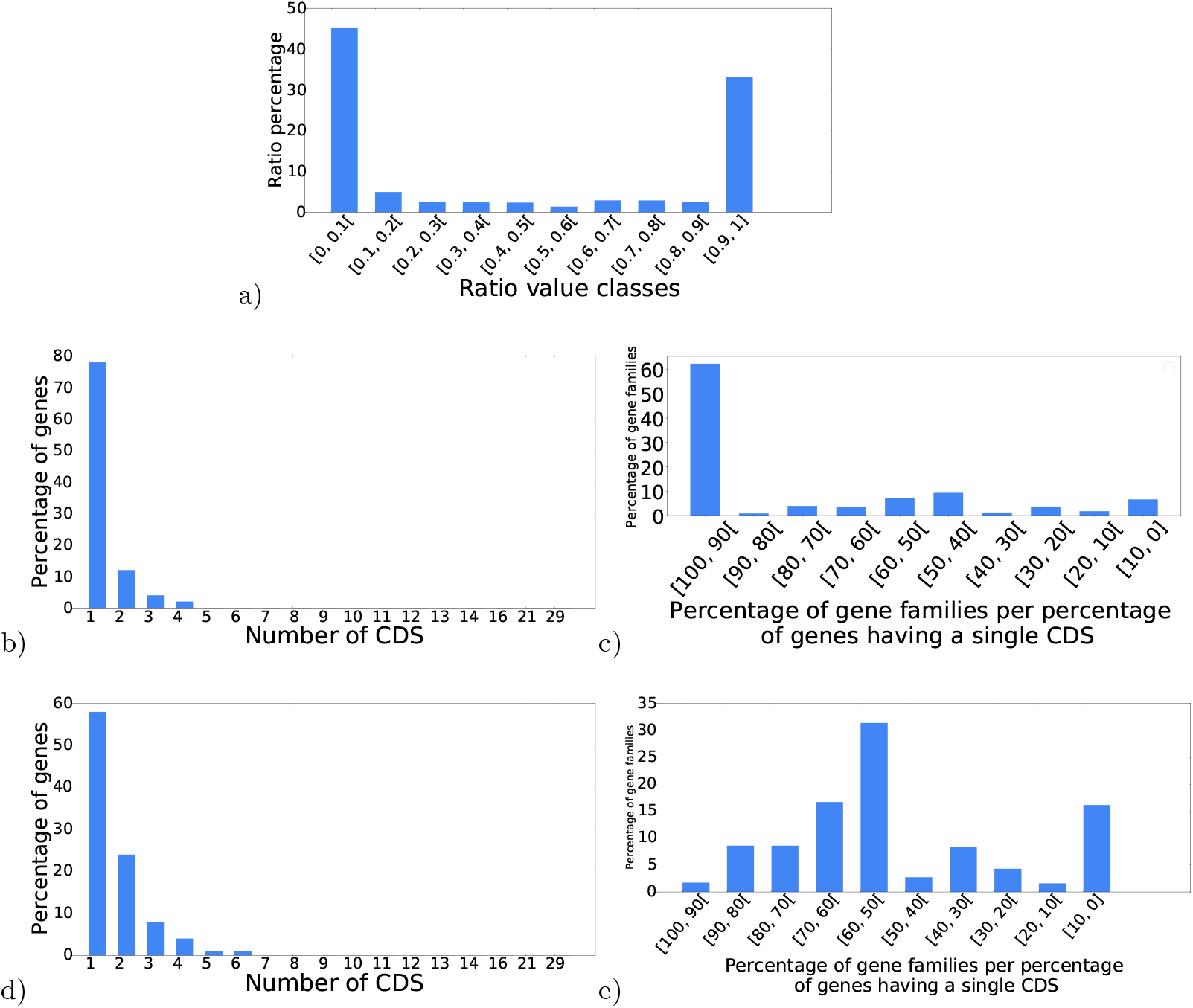
Statistics on the number of genes per family and the number of CDS per gene: **(a)** Percentage of gene families per ratio of the number of genes retained in this study over the number of genes in the EnsemblCompara database. **(b)** Percentage of genes per numbers of CDS in the 2036-dataset. **(c)** Percentage of gene families per percentage of genes having a single CDS in the 2036-dataset. **(d)** Percentage of genes per numbers of CDS in the 723-dataset. **(e)** Percentage of gene families per percentage of genes having a single CDS in the 723-dataset.

Despite the large number of single-CDS genes in the dataset, the comparative analyses between the LT, IsoSel and SHT reveal significant differences between the results of the two methods, as shown in the sequel.

The LT and IsoSel methods were applied on the gene families to reconstruct a set of gene trees respectively called the LT gene trees and IsoSel gene trees. Note that, for the same gene family, the LT gene tree may differ from the LT_db gene tree, because the LT_db is induced from the EnsemblCompara database gene tree that was often constructed on a larger set of genes as shown in Figure 5 (a). Using the SHT method, the gene trees were reconstructed for 7 different values of the parameter *α* of the SHT method, *α* = 0.0, 0.2, 0.4, 0.5, 0.6, 0.8, 1.0, which corresponds to the weight of the splicing-structure similarity in the similarity scores between CDS. This yielded 7 sets of gene trees for the SHT method called the SHT gene trees. For each value of *α*, the resulting SHT gene trees were compared with the corresponding LT, LT_db and IsoSel gene trees.

### 3.2 Comparison criteria

Four criteria are used to compare the SHT, IsoSel and LT gene trees for each value of *α* = 0.0, 0.2, 0.4, 0.5, 0.6, 0.8, 1.0. The first criterion is the percentage of common transcripts between the sets of representative transcripts computed in the LT and SHT method, and also between the sets of representative transcripts computed in the IsoSel and SHT method. This criterion is used to evaluate the similarity between the methods in terms of the set of selected representative transcripts. The second criterion is the percentage of trees from each method which pass an approximately unbiased (AU) test (Shimodaira 2002) used to evaluate the significance of a likelihood difference between candidate trees on the same set of taxa. The third criterion is the comparison of the reconciliation cost of the gene trees with the species trees. The last criterion is the comparison of species deduced from the set of gene trees built by each approach.

### 3.3 Comparison based on the percentage of common representative transcripts

Figure 6 shows the percentage of common representative transcripts between the LT and SHT methods. We observe that the representative transcripts chosen by the SHT method are not always the longest ones. Figure 6 (a) shows that for various values of *α*, the median percentage of common representative transcripts in gene families is almost 70% for the 723-dataset. Figure 6 (b) shows that when we consider only genes having more than one transcript, the median percentage of common representative transcripts decreases and it is almost 37%. Figure 6 (c) shows the average ratio of the mean similarity between the SHT transcripts over the mean similarity between the LT transcripts. As expected, we observe that the SHT transcripts are closer in terms of similarity than the LT transcripts. These results illustrate that the SHT method is effective at computing a set of representative transcripts different from the longest transcripts selected by the LT. The SHT representative transcripts are closer in terms of similarity than the LT transcripts are when we increase the weight *α* of the splicing structure similarity score.

**Fig. 6:**
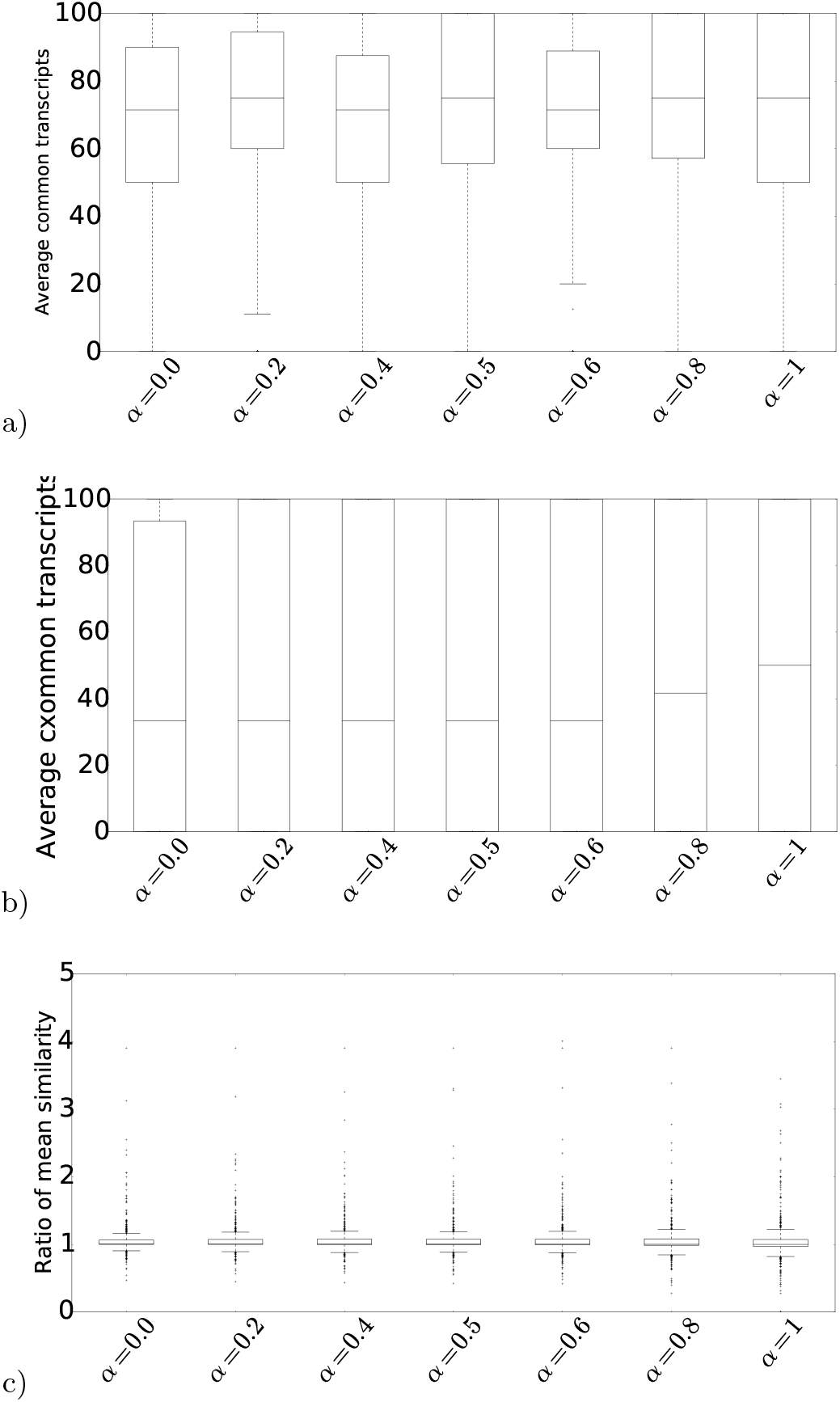
Common representative transcripts: Boxplot of percentages of common representative transcripts in gene families between the SHT and the LT methods per values of *α* **(a)** in the 723-dataset, **(b)** when considering only genes having more than one transcript in the 723-dataset. **(c)** Boxplot of ratios of the mean similarity between the SHT transcripts over the mean similarity between the LT transcripts.

We also observe very few changes between the results of various values of *α* that determine the weights of the splicing structure similarity and the sequence similarity scores in the CDS similarity measure. It suggests that there is a correlation between the splicing structure similarity and the sequence similarity scores. This is confirmed by the mean of the correlation scores between the two scores computed for all gene families of the 2036-dataset: minimum correlation score equals 0.22, maximum 1.0, mean 0.64, and median 0.57.

Figure 7 shows similar results when comparing the percentage of common representative transcripts between IsoSel and SHT methods. Figure 7 a) shows that for various values *α*, the median percentage of common representative transcripts in gene families is almost 70% for the 723-dataset. Figure 6 (b) shows that when we consider only genes having more that one transcript, the median percentage of common representative transcripts is almost 30%. Figure 7 c) shows that the SHT representative transcripts are closer in terms of similarity than the IsoSel representative transcripts.

**Fig. 7:**
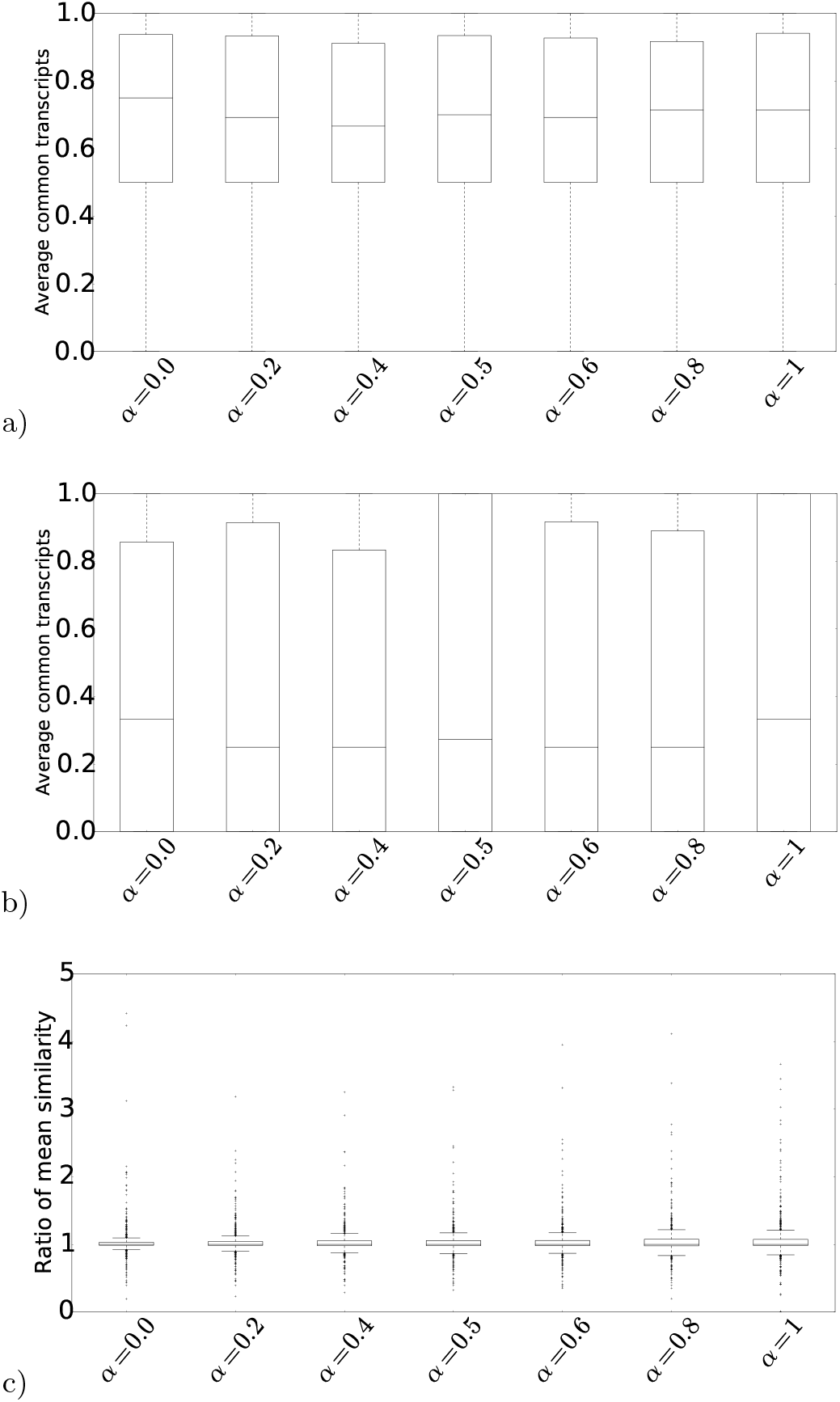
Common representative transcripts: Boxplot of percentages of common representative transcripts in gene families between the SHT and the IsoSel methods per values of *α* **(a)** in the 723-dataset, **(b)** when considering only genes having more than one transcript in the 723-dataset. **(c)** Boxplot of ratios of the mean similarity between the SHT transcripts over the mean similarity between the IsoSel transcripts.

### 3.4 Comparison based on the percentage of trees passing the AU test

We used the approximately unbiased (AU) test to compare the trees built by the LT and the SHT methods (ibid.). The AU test compares the likelihood of a set of trees in order to build a confidence set, which will consists of the trees that cannot be statistically rejected as a valid hypothesis. We say a tree can be rejected if its AU value, interpreted as a *p – value*, is under 0.05. Otherwise, no significant evidence allows us to reject one of the two trees. We start by executing PhyML (release 27) (Guindon and Gascuel 2003) on the trees to obtain the log-likelihood values per site and we execute Consel (release 28) to obtain the results from the AU test.

Figure 8 shows the percentage of LT_db, LT, IsoSel and SHT trees passing the AU test for various values of *α*. Since the AU test requires gene sequences, the sequences of transcripts selected by Ensembl, IsoSel and SHT have been used as gene sequences. Figures 8 a), b), and, c) respectively illustrates the results obtained when the gene sequences are the transcript sequences selected by SHT, LT, and, IsoSel. When the transcript selected by SHT are used, we clearly observed that the SHT method slightly better than LT_db, LT and the IsoSel method. Between 78% to 79% of the SHT gene trees pass the AU test. 79% of trees passing the AU test are obtained for *α* = 0.8. Whereas, for the LT_db trees, this percentage varies between 68% and 70%, for LT trees, this percentage varies between 74% and 75%, and for IsoSel trees between 76% and 77%. When transcripts selected by LT Figure 8 b) or IsoSel 8 C) are used, we observe that each both methods performs also slightly better. This can be explain by the fact the transcript sequences choose for AU test favors the the tree having the transcript better than other trees. For this criteria, the results clearly illustrate that the gene trees from the fourth approaches are all comparable based on the results of AU test. The SHT and IsoSel gene trees also allow to identify cases in which the LT gene trees do not pass the AU test, this confirms that the choice of the longest transcript is not always the best approach to select a set of representative transcripts. The results also allow to compare the accuracy of the LT and LT_db gene trees. They show that using a larger set of genes provides higher accuracy for the LT_db gene trees as compared to the LT trees that are computed on a restricted set of genes.

**Fig. 8:**
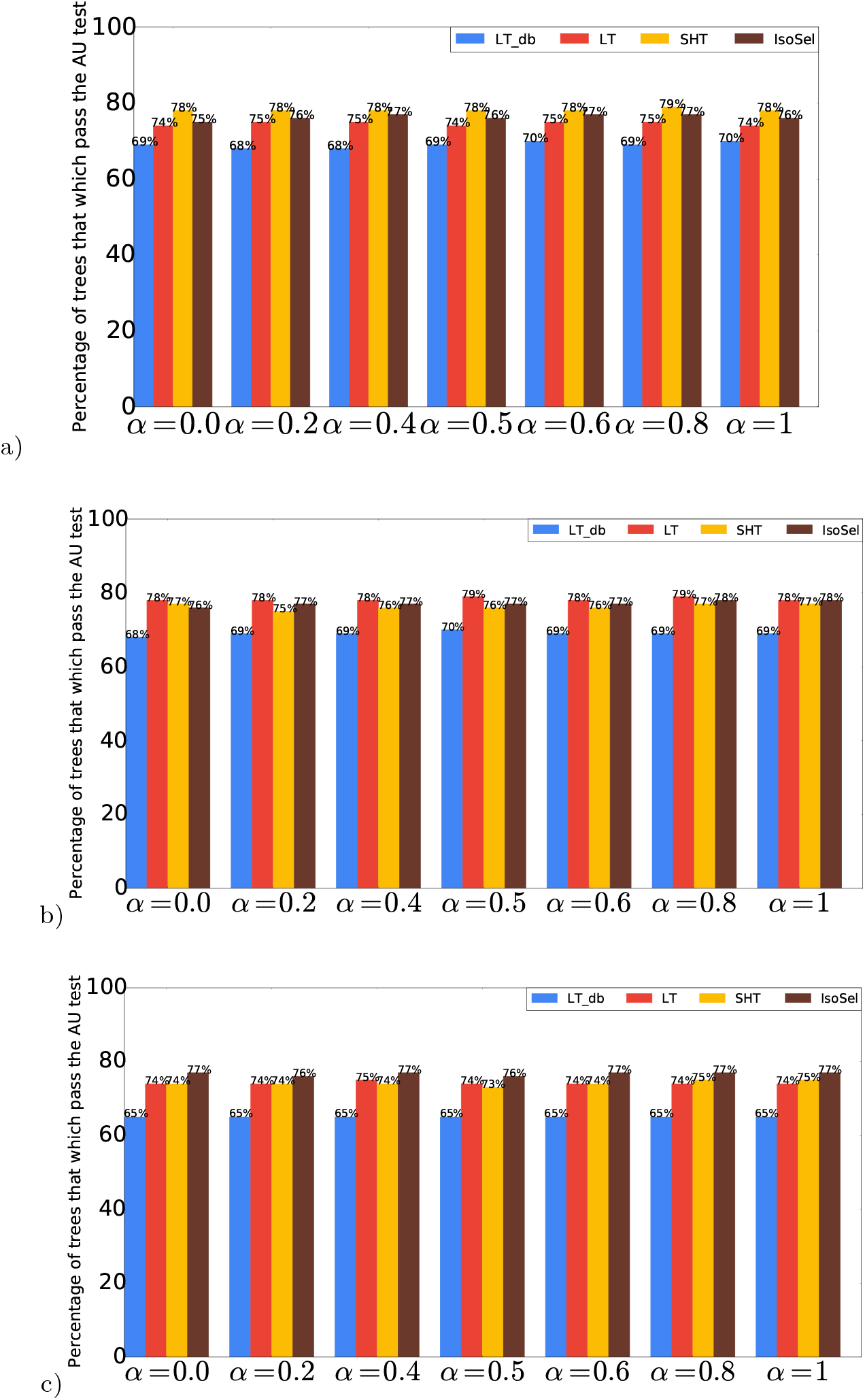
AU test: Percentage of trees from each method (LT_db, LT, SHT, IsoSel) which pass the AU test per values of *α*. **(a)** When the transcript sequences selected by SHT are used as gene sequences for AU test. **(b)** Same comparison when transcript sequences selected by LT are used. **(c)** Same comparison when transcript sequences selected by IsoSel are used.

### 3.5 Comparison based on the reconciliation cost with a species tree

Figure 9 (a) shows the comparative results between the SHT, and, LT gene trees in terms of reconciliation cost on the 723-dataset. We observe that, while 65% of gene tree build by SHT and LT are eqauls, 20% are differents but have the same reconciliation cost. For the remaining gene trees, SHT always have a highest percentage of tree having the lowest reconciliation cost when varying the values of *α*. This percentage varying between 8 to 10%, while the same percentage for LT varying between 6 to 8%. Figure 9 (b) shows the comparative results between the SHT and the LT_db gene trees on the 2036-dataset. Here, 72% of gene tree build by both approaches are equals, while almost 17% are differents but have the same reconciliation cost. For the remaining gene trees, SHT always have the highest percentages of gene trees having the lowest reconciliation cost when varying the values of *α* Figure 9 (c) shows the comparative results between the SHT and the IsoSel gene trees on the 2036-dataset. Here, the percentages of equal gene trees build by SHT and IsoSel is almost 87%. While the percentage of differents gene trees having the same reconciliation cost varying between 8 to 10%. For the remaining gene tree, there is remarkable difference, while only 1% of gene tree build by IsoSel have a lowest reconciliation cost, 3 to 4% of gene trees build by SHT have lowest reconciliation cost. This results supports the use of the new similarity measure and clustering method to select sets of representative transcripts more wisely.

**Fig. 9:**
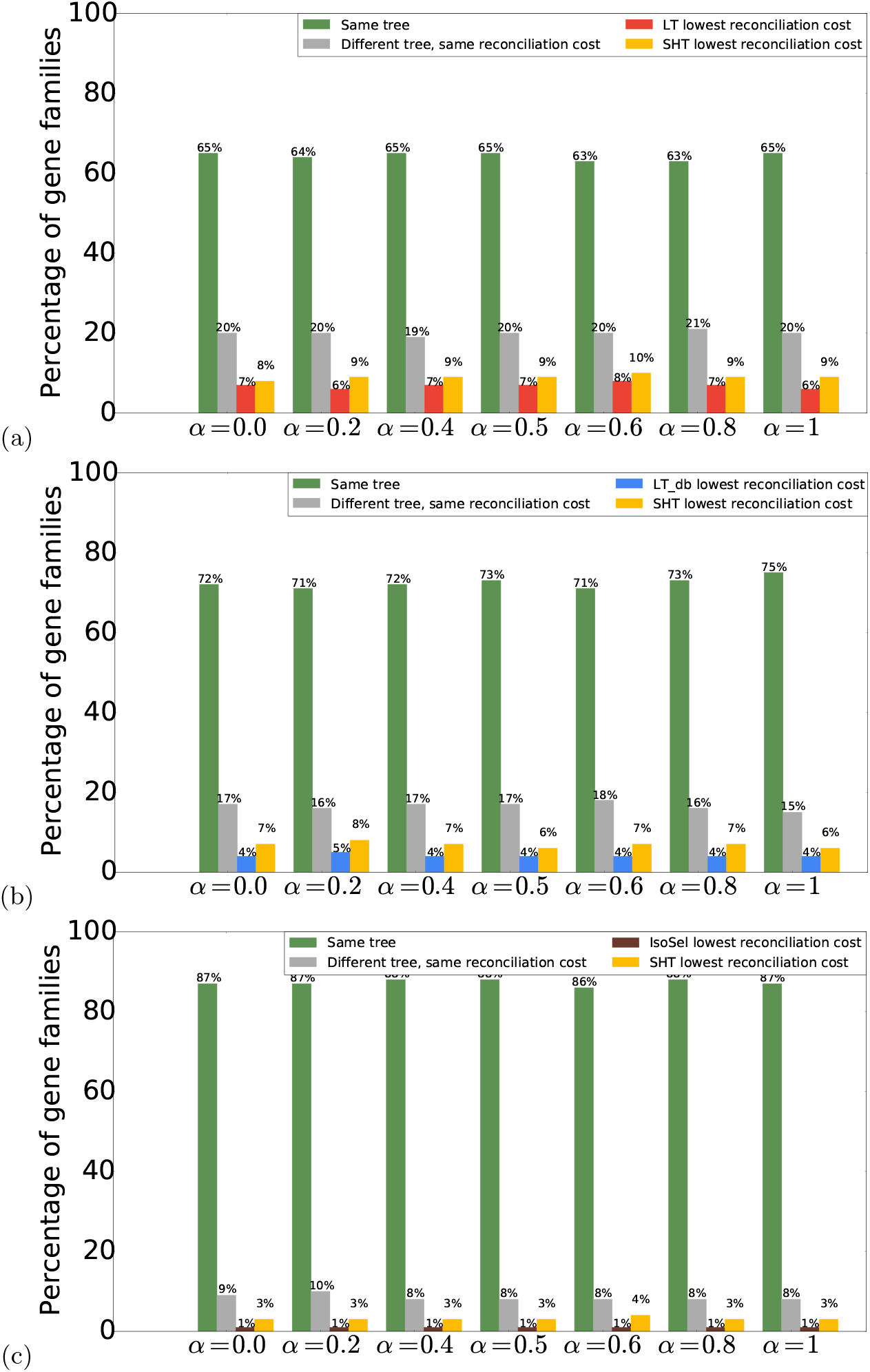
Reconciliation cost: **(a)** Percentage of gene families for which the SHT and the LT gene trees are identical (green), for which the SHT and the LT gene trees differ but that they have the same reconciliation cost with the species tree (grey), for which the LT gene tree has a lower reconciliation cost (red), and for which the SHT gene tree has a lower reconciliation cost (yellow), per values of *α*. **(b)** Same for the comparison between the SHT and the LT_db gene trees. **(c)** Same for the comparison between the SHT and the IsoSel gene trees.

This result, coupled with the results of Figure 6 and 7 on the fourth methods’ accuracy comparison, (LT *<* LT_db *<* IsoSel *<* SHT) shows that choosing wisely the set of representative transcripts (i.e SHT trees) allows to reach the accuracy obtained using a larger set of genes, and even to compute more accurate gene trees.

### 3.6 Comparison based on the evaluation of the deduced species trees

The set of gene trees built by LT, LT_db, IsoSel, and, SHT (*α* = {0.0, 0.2, 0.4, 0.5, 0.6, 0.8, 1.0}) were used as input of Astral (Mirarab and Warnow 2015) to build a species tree. 5 percentages (20%, 40%, 60%, 80%, and, 100%) of gene trees were used as input of Astral. We used criteria two criteria to compare theses species trees. The first criterion is the quartet support values of branches of each species. The quartet score is the proportion of input gene tree quartet trees satisfied by the species tree. This is a number between zero and one; the higher this number, the less discordant your gene trees are. Figure 10 shows the comparison results between LT, LT_db, IsoSel, and the 7 species trees obtained each value of *α*. We note on this boxplot that the quartet support values of each method increase with the percentage of gene trees used. When 20%, 40,%, 60% or 80% of gene trees are used, the quartet support values of methods are slightly equals. When 100% of gene trees are used, the quartet support values of SHT is higher compare to one of LT, LT_db, and, SHT. Figure 11 shows the percentages of common clades between reconstruct species trees when varying the percentage of gene trees used and the initial Ensembl species tree. We observe that SHT has the highest percentage of common clades with the species tree.

**Fig. 10:**
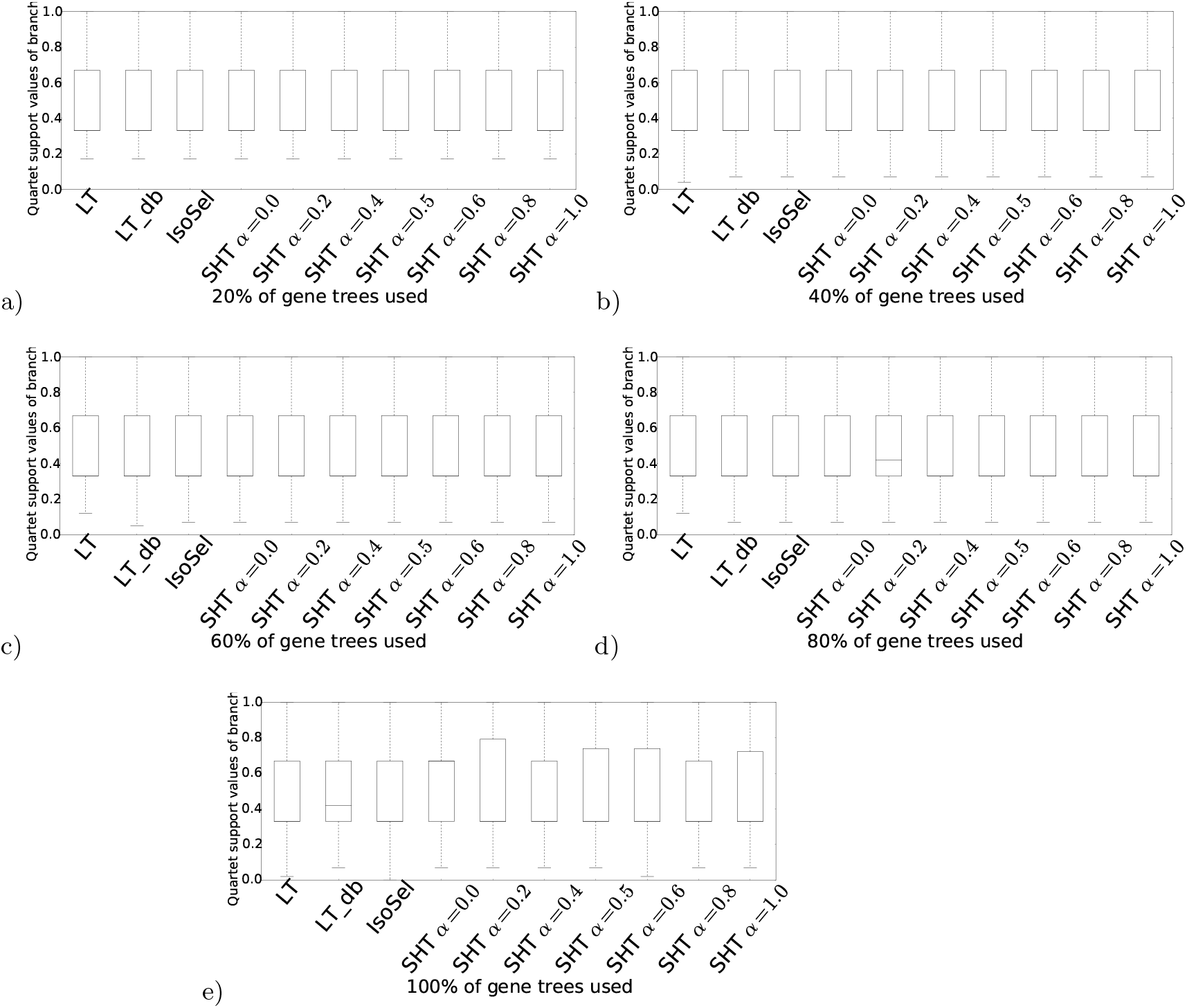
Species trees: Quartet support values of branches for species tree deduced from the set of gene trees build by each of approach LT_db, LT, IsoSel, SHT. **(a)** When 20% of gene trees are used as input of Astral to build species tree. **(b)** Same when 40% of gene are used. **(c)** Same when 60% of gene are used. **(d)** Same when 80% of gene are used. **(e)** Same when 100% of gene are used.

**Fig. 11:**
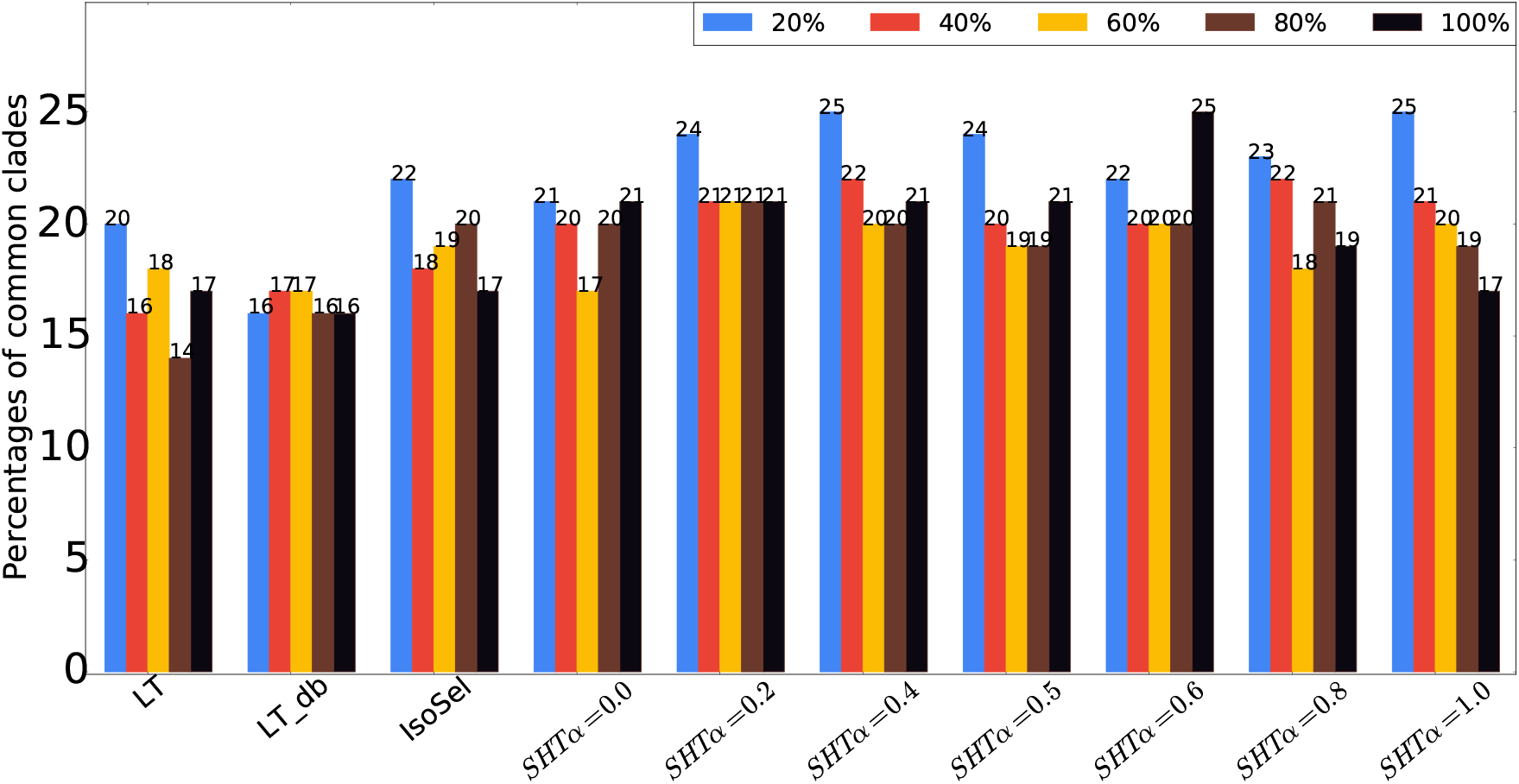
Percentages of common clades between each reconstruct species trees and the initial Ensembl species tree when varying the percentage of used gene trees from.

## 4 Conclusion

A new approach called Splicing Homology Transcript (SHT) method has been devised to estimate eukaryote gene trees. It is based on a novel and alternative way to choose the representative transcripts of a set of homologous genes. Contrary to the most used approaches that select the longest transcript of each gene as the representative transcript independently of other homologous genes, it selects a set of representative transcripts using a combination of the sequence similarity and the splicing structure similarity between sequences. This approach yields more accurate gene trees that the approach that selects the longest transcript of each gene as the representative transcript. We predict that alternative-splicing-aware gene tree reconstruction methods will be instrumental to advancing our understanding of gene family and genome evolution.

The analysis of the sequence and the structure similarity scores between transcripts of a gene family shows that the two scores are correlated. In this study, we have enriched the similarity measure definition based on the sequence by including information related to the structure, it will be interesting as a future work to carry out machine learning in order to determine the weight allowing to obtain better results. As long as this study exploit the spliced structure of sequences, the use of well-annotated species may increase the results accuracy. Moreover, the SHT method differs from the EnsemblCompara and IsoSel gene tree estimation method only by the strategy used in the representative transcripts’s selection step. Future works will consider alternative strategies in the other steps, such as the multiple sequence alignment and the phylogeny estimation steps. The method also selects a single set of representative transcripts while there can be several co-optimal sets of representative transcripts. Future extensions of the method will include the computation of a transcript tree for each co-optimal set of representative transcripts, and the combination of the resulting trees to increase the precision of the final gene tree. Finally, a further extension of this work will consist of combining the set of representative transcript trees into a transcript phylogeny representing the evolutionary history of all transcripts of a set of homologous genes.

## 5 Acknowledgements

The authors thank the CoBiUS lab at University of Sherbrooke for their helpful constructive discussions.

## Notes

### Competing Interest Statement

The authors have declared no competing interest.

